# Polyamines Mediate Folding of Primordial Hyper-Acidic Helical Proteins

**DOI:** 10.1101/2020.10.01.321877

**Authors:** Dragana Despotović, Liam M. Longo, Einav Aharon, Amit Kahana, Tali Scherf, Ita Gruic-Sovulj, Dan S. Tawfik

**Author notes:** These authors contributed equally to this work (Dragana Despotović, Liam M. Longo). Corresponding authors: E-mail addresses.

## Abstract

Polyamines are known to mediate diverse biological processes, and specifically to bind and stabilize compact conformations of nucleic acids, acting as chemical chaperones that promote folding by offsetting the repulsive negative charges of the phosphodiester backbone. However, whether and how polyamines modulate the structure and function of proteins remains unclear. Further, early proteins are thought to have been highly acidic, like nucleic acids, due to a scarcity of basic amino acids in the prebiotic context. Perhaps polyamines, the abiotic synthesis of which is simple, could have served as chemical chaperones for such primordial proteins? We replaced all lysines of an ancestral 60-residue helix-bundle protein to glutamate, resulting in a disordered protein with 21 glutamates in total. Polyamines efficiently induce folding of this hyper-acidic protein at sub-millimolar concentrations, and their potency scaled with the number of amine groups. Compared to cations, polyamines were several orders of magnitude more potent than Na^+^, while Mg^2+^ and Ca^2+^ had an effect similar to a di-amine, inducing folding at approximately seawater concentrations. We propose that (*i*) polyamines and dications may have had a role in promoting folding of early proteins devoid of basic residues, and that (*ii*) coil-helix transitions could be the basis of polyamine regulation in contemporary proteins.

## Introduction

Natural polyamines contain two or more amino groups separated by aliphatic hydrocarbon chains, and can be either linear or branched. At neutral pH, polyamines are polycations that – unlike metal ions, which are effectively point charges – encode patterns of regularly spaced positively charged sites. Low molecular weight, linear polyamines (see **Figure 1** for the most common natural examples) are present in all living organisms (1) and essential for cell survival (2). The functional profile of these and other polyamines is diverse, and these simple metabolites mediate a variety of biological processes, including chromatin structure remodeling (3), gene transcription and translation (4, 5), cell proliferation (6), and circadian clock regulation (7). Although the intracellular concentrations of polyamines are not well characterized, current evidence suggests it can be quite high. The concentrations of spermidine and putrescine in *E. coli*, for example, are estimated to be about 6 mM and 20 mM, respectively (8), though the fraction of bound versus free polyamine is unknown. Intracellular polyamine concentrations have also been shown to vary significantly with cellular state, thus suggesting a regulatory role for polyamines (9). Mechanistically, polyamines operate on nucleic acids through interactions with the negatively charged phosphodiester backbone resulting in changes in DNA conformation (10, 11), tRNA stabilization (12), and even phase separation (13). In the laboratory, polyamines have been employed to promote folding and stability of nucleic acids *in vitro* (14–16). However, with respect to proteins, modulation of structure and/or function by polyamines has centered largely on amyloidogenesis (17–19); consequently, the mechanisms by which protein structure-function can be modulated by polyamines are largely unknown.

**Figure 1.**
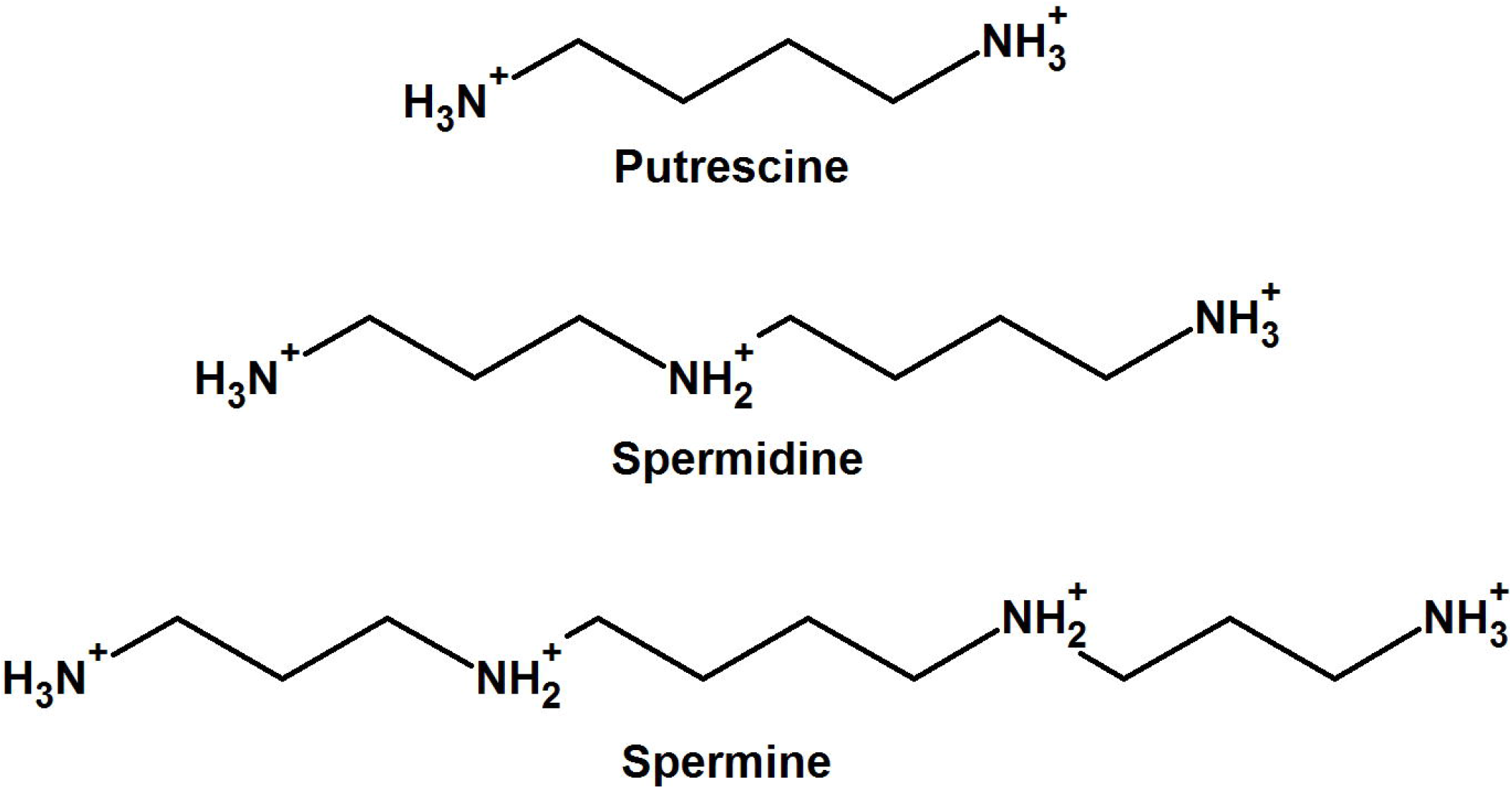
Natural polyamines used in this study. Polyamines are polycations at neutral pH, with p*K*_a_s that range from 8-10.9 (50, 51).

A second interesting aspect of polyamines is their potential role at the dawn of life. Given the ubiquity, essentiality, and relatively simple chemical structure of polyamines, it is likely that they are ancient metabolites that emerged early in life’s history, perhaps as chemical chaperones for early nucleic acids, including ancient ribozymes (20). Consistent with this view, primordial synthesis of polyamines has been observed (21). Here, we ask whether polyamines might have had a role in the emergence of the early proteins. Modern proteins are comprised of both positively and negatively charged amino acids, and the charge of most proteins is either moderately negative or moderately positive to maximize foldability and solubility at around neutral pH. While solubility may be enhanced by significant negative charge, foldability will be hindered. In the absence of positively-charged residues, salt bridges – which are critical for protein stability, as reflected in thermophilic proteins (22) – are lacking, leaving only the repulsive interactions between adjacent negative charges that destabilize compact structures. This may not be an issue in stable, modern proteins (23), but is a severely limiting factor in the absence of a large well-packed hydrophobic core, as expected at the early stages of protein evolution. In early life contexts, the availability of the contemporary basic amino acids was likely poor (24, 25). Alternative basic amino acids such ornithine could have been a substitute (26, 27), but the possibility that basic amino acids were depleted in, or even entirely absent from, the first proteins cannot be ruled out (28–30). Thus, understanding whether and how highly acidic proteins could fold or function remains a major unsolved problem in early protein evolution. Furthermore, folding of acidic proteins is fundamentally the same problem faced by the ribozyme, which employs chemical chaperones, most frequently dications, to enable globular packing. Perhaps the earliest proteins were similarly dependent on chemical chaperones?

To determine if polyamines can operate as efficient chemical chaperones for acidic proteins, we constructed a hyper-acidic version of a previously described reconstructed ancestral protein comprising a tandem duplication of a helix-hairpin-helix (HhH) motif (**Figure 2**) (26, 31). This protein, dubbed Acidic-(HhH)_2_, is devoid of arginine and lysine and has 21 glutamate residues out of 60 residues in total. As expected, Acidic-(HhH)_2_, is unfolded in low salt buffer and required near-molar concentrations of NaCl to achieve complete folding. Addition of submillimolar concentrations of polyamines, however, induced formation of α-helical structure. Our results inspire the hypothesis that polyamines served as chemical chaperones of ancient proteins, and that this feature may exist in polyamine-regulated proteins today.

**Figure 2.**
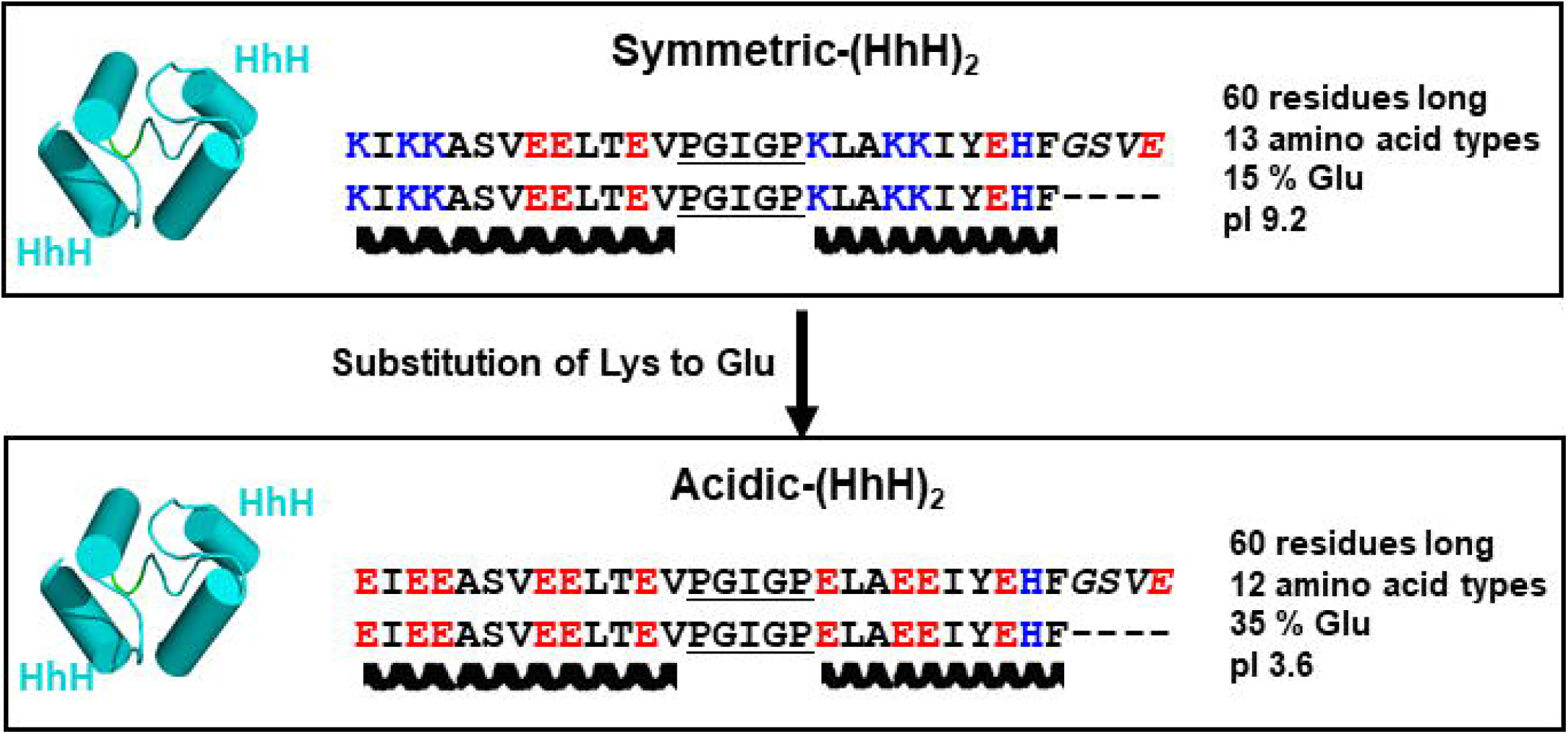
Design of Acidic-(HhH)_2_. Symmetric-(HhH)_2_ is a fully symmetric protein constructed in a previous study to understand the properties of ancient protein forms (26). Derived from a family of dsDNA binding proteins, it is positively charged at neutral pH. To generate a hyper-acidic model protein for this study, all lysine residues in Symmetric-(HhH)_2_ were substituted to glutamate (see *Results* for more details). The conserved loop residues between the two helices of the HhH motif (PGIGP) are underlined and the linker residues (GSVE) between the two HhH motifs are rendered in italics. The C-terminus of Acidic-(HhH)_2_ bears a 6x His-tag connected by a two-residue Leu-Glu linker (not shown).

## Materials and Methods

### Protein Expression and Purification

Synthetic genes were ordered from Twist Bioscience (www.twistbioscience.com) and cloned into a pET21a vector, yielding variants with a C-terminal 6xHis tag. All constructs were verified by Sanger sequencing. Transformed *E. coli* BL21(DE3) cells were induced at OD_600_ = ~0.6 with 1 mM IPTG. Induced cells were shake-incubated at 20 °C overnight. Cell pellets were collected by centrifugation and frozen at −20 °C for storage. Cell pellets from 250 mL cultures were resuspended in 30 mL of 100 mM NaCl, 50 mM Tris/HCl pH 7.5, 2.5 U/ml Benzonase (Merck Millipore), 0.2 U/ml rDNase1 (Thermo Fisher Scientific), 0.25x Protease Inhibitor Cocktail for Histidine-tagged proteins (Sigma-Aldrich), and 0.3 mg/mL lysozyme, and shake-incubated at 37 °C for 90 minutes. The lysates were then cooled on ice for 15 minutes and sonicated for 5 minutes, with a 30 s cool-down period after every 30 s of sonication. After cell lysis, the samples were spiked with 0.5 M NaCl and 5 mM imidazole and gently rocked for 30 min at room temperature (moderate concentrations of NaCl decreased unspecific binding to Ni-NTA and also promoted solubility of the parent protein (26)). Pellets were clarified by centrifugation and passed through a 0.45 μm sterile filter. Clarified lysates were applied to 3 mL packed Ni-NTA resin pre-equilibrated in 5 mM imidazole, 0.5 M NaCl, 50 mM Tris/HCl, pH 8.0 (“purification buffer”). After sample application, the resin was washed with 25 mL of purification buffer. Nonspecifically bound proteins were removed by washing with purification buffer spiked with 25 mM imidazole. The bound protein was eluted with 20 mL of 500 mM imidazole, 50 mM Tris/HCl, 500 mM NaCl. In samples used for NMR analysis, an additional wash step of 20 mL 6 M GuHCl was performed. The eluted protein fractions were dialyzed against 20 mM bis-Tris pH 5.5, 50 mM NaCl using a 3.5 kDa MWCO SnakeSkin Dialysis Tube (Thermo Fisher Scientific, Waltham, Massachusetts). Next, the dialyzed sample was loaded on an anion exchange HiTrap Q HP 5 mL column (GE healthcare life sciences, Boston, Massachusetts) pre-equilibrated with 20 mM bis-Tris pH 5.5, 50 mM NaCl at a flow rate of 1 mL/min using an FPLC AKTA-prime plus chromatography system (GE, Boston, Massachusetts). The protein was eluted by a linear gradient of NaCl (0.05-1 M NaCl in 20 mM bis-Tris pH 5.5) over the course of 20 column volumes at flow rate 2 mL/min. The fractions containing Acidic-(HhH)_2_ were combined and dialyzed against 5 mM Tris pH 7.5, 25 mM NaCl. Finaly, the sample was concentrated using a 3 kDa MWCO centrifugal filtration unit (Thermo Fisher Scientific) and protein concentration was measured with a BCA kit (Thermo Fisher Scientific).

### Circular Dichroism (CD) Spectroscopy

CD spectra were collected on a Chirascan circular dichroism spectrometer (Applied Photophysics). Samples containing 5 μM protein in 5 mM Tris/HCl, 25 mM NaCl, pH 7.5 (unless otherwise stated) were placed in a 1 mm pathlength quartz cuvette and equilibrated at 25 °C. Tris buffer, although not ideal for CD studies, was chosen because phosphate buffer precipitated in the presence of polyamines. Spectra were collected at the range of 195-260 nm with a data pitch of 1 nm and a slit width of 1.5 nm. The data points exceeding 700 V applied to the photomultiplier tube were discarded. Titrations were performed by sequential additions of stock polyamine solutions or salt solutions in 5 mM Tris/HCl, 25 mM NaCl adjusted to pH 7.5. The reported spectra were buffer-subtracted and corrected for dilution caused by added titrant.

### Nuclear Magnetic Resonance (NMR) Spectroscopy

^1^H NMR spectra were recorded for 100 μM Acidic-(HhH)_2_ in 25 mM fully deuterated Tris (Tris-d11) pH 7.5, 25 mM NaCl, in 90% H_2_O/10% D_2_O and upon addition of various spermine concentrations. Titrations were performed by sequential additions of a pH-adjusted, 1 M spermine solution in 25 mM NaCl, 90% H_2_O/10% D_2_O. NMR experiments were conducted at 293 K on a Bruker AVANCE NEO 600 MHz NMR spectrometer equipped with 5-mm cryogenic triple-resonance HCN TCI probe (triple axis X,Y,Z-gradients). Data were processed and analyzed using TOPSPIN 4.0 (Bruker BioSpin, Germany). The one-dimensional ^1^H NMR spectrum was acquired using excitation sculpting (32) to suppress the solvent signal; the 2D TOCSY spectrum (total correlation spectroscopy, (33)) at the spermine saturating concentration (1:250 molar ratio of protein:spermine) was recorded using a dipsi2 mixing time of 120 ms.

## Results

### Design of Acidic-(HhH)_2_

To determine if polyamines can induce folding of a primordial, hyper-acidic protein, we designed Acidic-(HhH)_2_ (**Figure 2**) based on our recently published Symmetric-(HhH)_2_ protein (26). Briefly, Symmetric-(HhH)_2_ was the result of ancestral inference and targeted simplification of the ancient and widely-distributed (HhH)_2_ protein fold. The (HhH)_2_ protein fold is an α-helix bundle formed by two symmetrically juxtaposed helix-hairpin-helix (HhH) motifs (34). The HhH motif is a pre-LUCA structure element (31) that interacts with nucleic acids. It is part of numerous proteins, including ribosomal proteins and polymerases, but when duplicated can form a stand-alone domain. The sequence of Symmetric-(HhH)_2_ is derived from the symmetrization of the reconstructed ancestor of all known (HhH)_2_ protein families. As the name implies, the sequence of the first and second HhH subdomains of Symmetric-(HhH)_2_ are identical. Symmetric-(HhH)_2_ therefore represents an intermediate along the trajectory leading from a primordial single HhH polypeptide to the contemporary (HhH)_2_ protein domain.

To generate Acidic-(HhH)_2_, all of the lysine residue in the Symmetric-(HhH)_2_ were mutated to glutamate, an abiotic, early-emerging amino acid (30). Glutamate was chosen over aspartate because it has a higher α-helix propensity (35) and because this mutation is preferred in the BLOSSOM62 substitution matrix (36). Twelve positions were exchanged in Symmetric-(HhH)_2_ to give Acidic-(HhH)_2_. The resulting protein is 60 residues long, has 100% sequence identity between the two HhH domains, and is comprised of just 12 amino acid types. In total, Acidic-(HhH)_2_ has 21 glutamate residues, no arginine or lysine, and just two histidine residues. Consequently, Acidic-(HhH)_2_ has a pI of 3.6 and bears a significant negative charge at neutral pH. In buffer at pH 7.5, Acidic-(HhH)_2_ is unfolded, as demonstrated by random coil signal in the CD spectrum (**Figure 3**) and the poor peak dispersion in the 1D ^1^H NMR spectrum (**Figure 4**, top).

**Figure 3.**
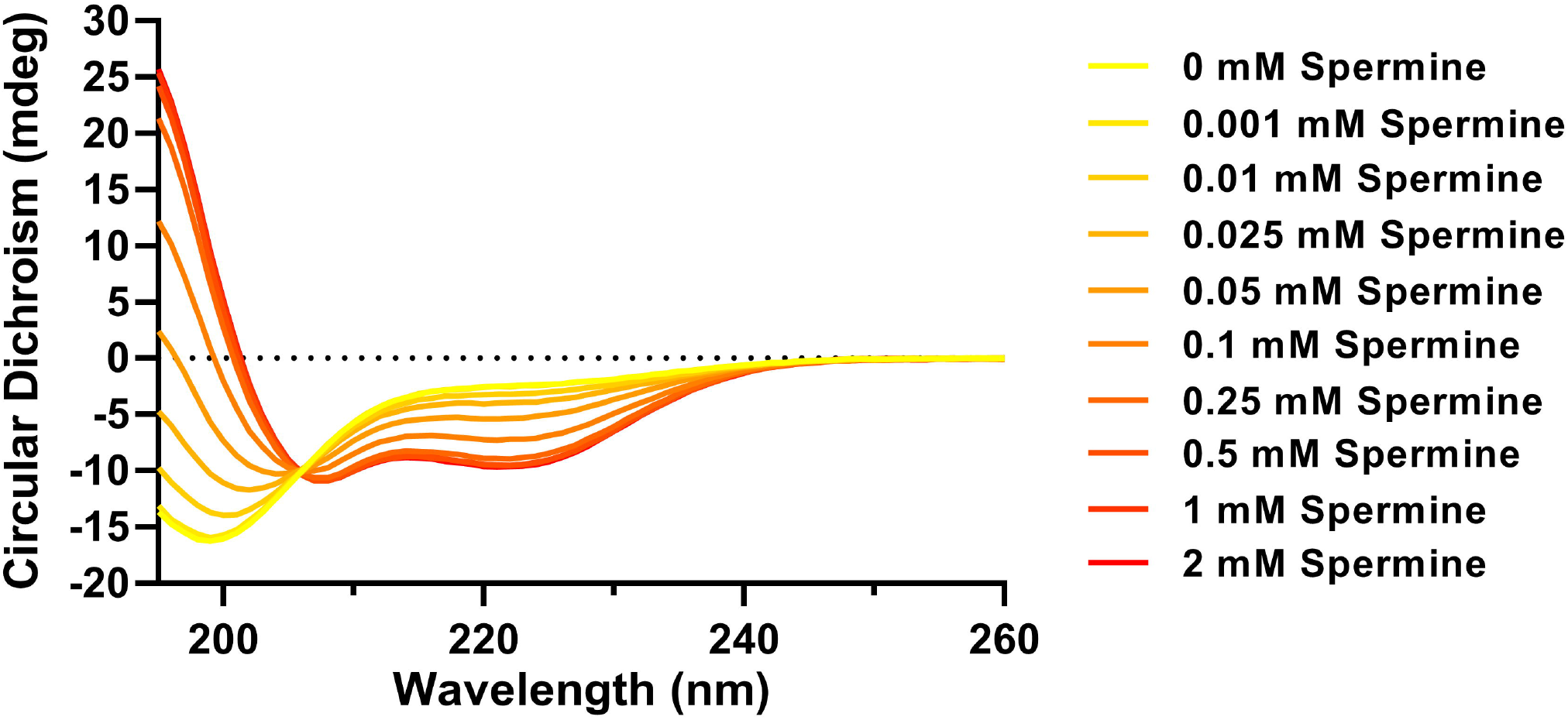
Folding of Acidic-(HhH)_2_ upon spermine addition monitored by circular dichroism (CD). Shown are CD spectra of 5 μM Acidic-(HhH)_2_ with spermine at varying concentrations. Each curve represents the average of two scans after buffer subtraction (5 mM Tris pH 7.5, 25 mM NaCl) and correction for dilution by added titrant. The development of a peak at <195 nm, and negative bands at ~ 208 nm and ~222 nm suggests folding into a predominantly α-helical conformation. An isodichroic point at ~206 nm suggests a two-state transition.

**Figure 4.**
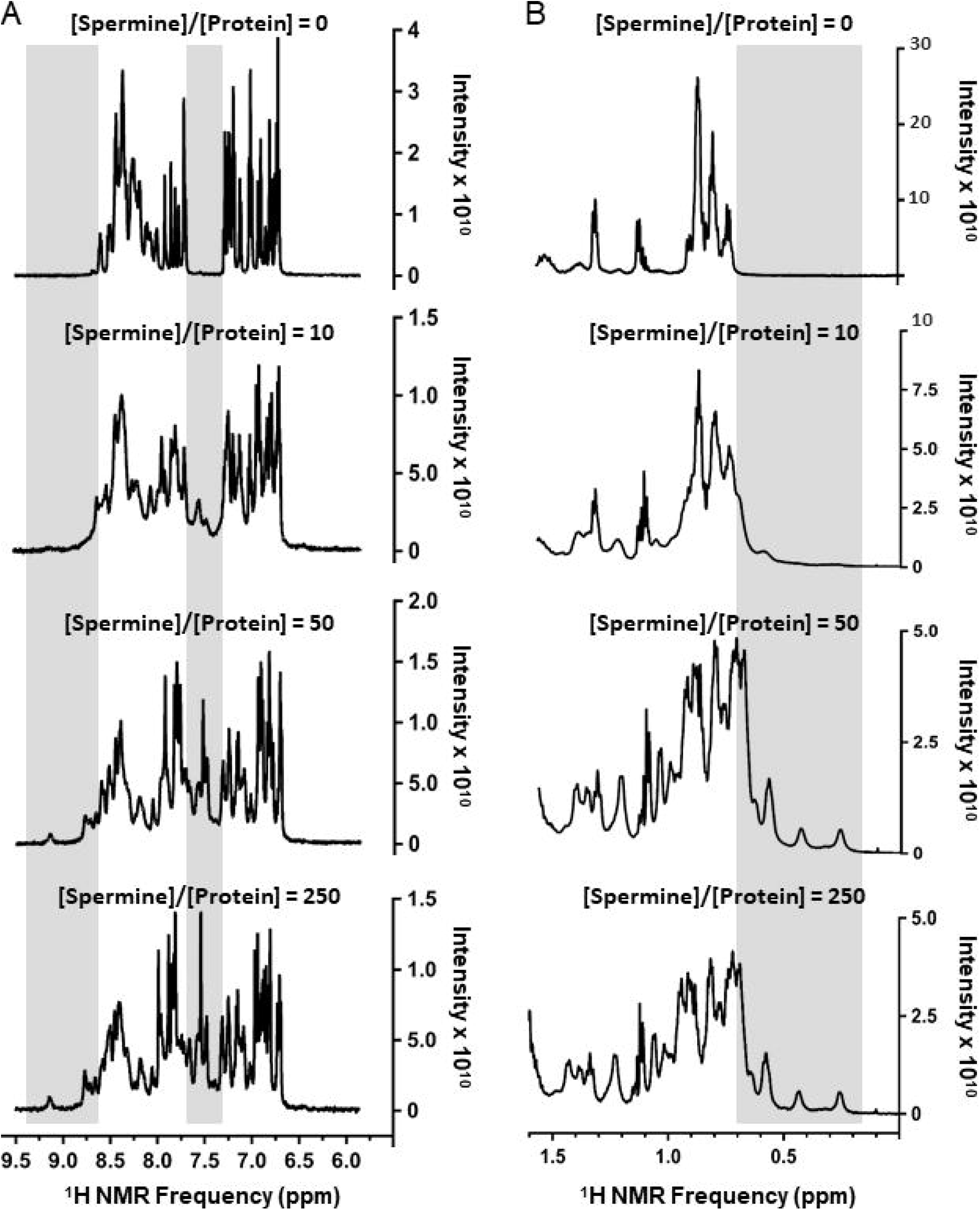
Titration of Acidic-(HhH)_2_ with spermine monitored by NMR. 1D ^1^H NMR spectra of 100 μM Acidic-(HhH)_2_ in varying concentrations of spermine (0, 1, 5 and 25 mM). The increasing peak dispersion, and observation of new peaks upon addition of spermine (highlighted with a gray background), both indicate that spermine is promoting the acquisition of structure. (A) Zoom-in on the spectral region reporting amide and aromatic protons; (B) Spectral region corresponding to aliphatic protons, predominantly methyl groups.

### Spermine induces α-helix formation

The effect of spermine, a natural polyamine with 4 amino groups separated by alkyl chains (**Figure 1**), on the structure of Acidic-(HhH)_2_ was first monitored by CD spectroscopy. Upon addition of sub-millimolar concentrations of spermine, signals associated with α-helical structure rapidly developed in the CD spectrum during the deadtime of the experiment (**Figure 3**; **Figure S1**). Titration of Acidic-(HhH)_2_ with spermine yielded a midpoint of the folding transition of ~70 μM (**Figure 5**, see legend). The spectra from the spermine titration also revealed the presence of an isodichroic point at around 206 nm, consistent with a simple two-state transition between a random coil conformation and an α-helix conformation. The low concentration at which we observed the folding transition indicates that the effect of spermine is not simply from screening negative charge: The circular dichroism buffer includes 25 mM NaCl (ionic strength = 25 mM), and addition of 0.1 mM of spermine increased the ionic strength by just 1 mM (assuming spermine is completely protonated and neutralized by chloride ions).

**Figure 5.**
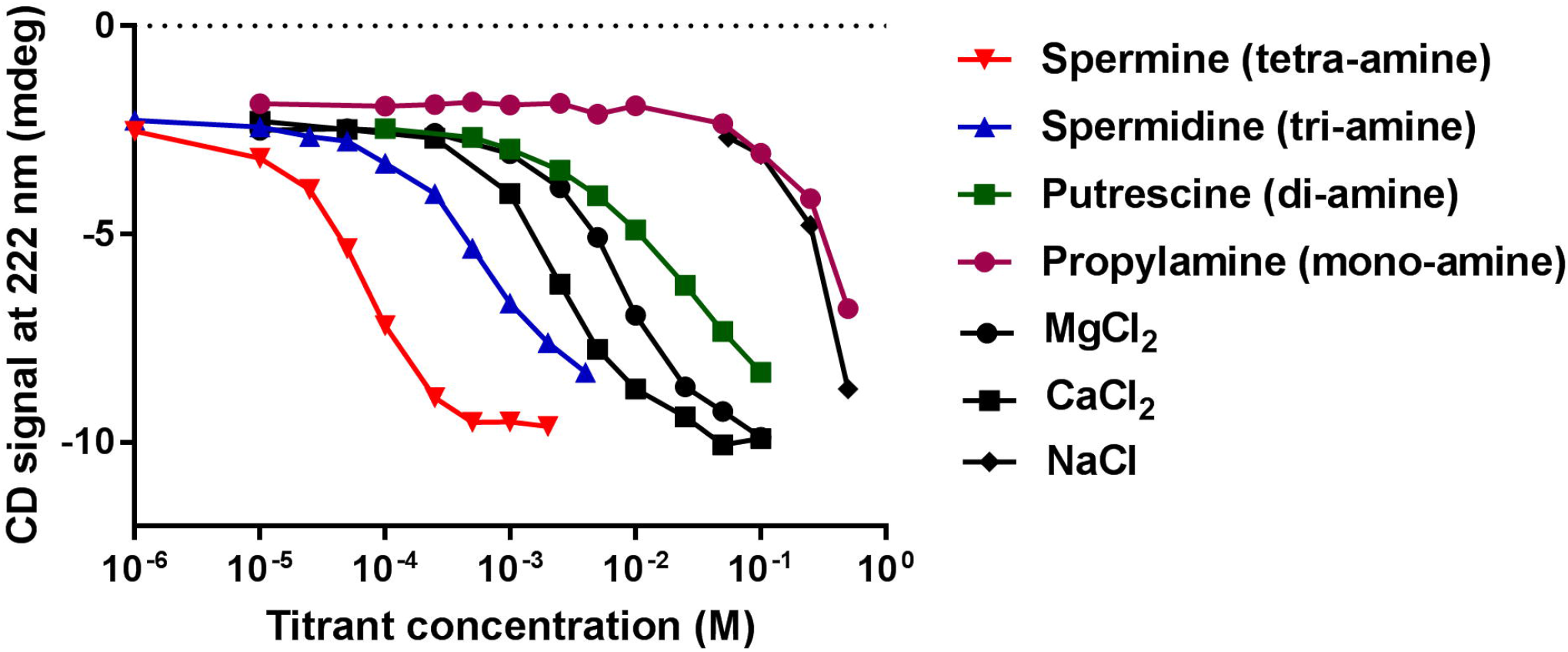
Titration of Acidic-(HhH)_2_ with various polyamines and salts. Circular dichroism spectra of 5 μM Acidic-(HhH)_2_ upon addition of various polyamines and salts were collected (**Figure S4**). Plotted here is the CD signal at 222 nm, a reporter of α-helical structure. Empirical midpoint concentrations are: spermine = 0.07 mM, spermidine = 0.7 mM, putrescine = 21 mM, propylamine = 420 mM, CaCl_2_ = 2.6 mM, MgCl_2_ = 7.4 mM, and NaCl = 300 mM. The midpoint concentrations were estimated from a linear interpolation between points and assuming a folded signal of approximately −10 mdeg at 222 nm. Instead of a technical duplicate, we performed a second polyamine titration experiment with an independent protein preparation at twice the protein concentration, presented in **Figure S3**. The midpoint concentrations between the two titrations are within 20%, and the order of potency is the same.

To cross validate our results from circular dichroism, the titration of Acidic-(HhH)_2_ with spermine was followed by ^1^H NMR. In the absence of spermine, Acidic-(HhH)_2_ exhibits poor chemical shift dispersion (**Figure 4**, top). This observation is indicative of an unfolded state, as the dispersion of the ^1^H resonances, and specifically of the ^1^HN, ^1^H and methyl signals, is much greater in folded proteins, reflecting the well-known dependence of the environment of proton nuclei on secondary and tertiary structure (37). Titration of Acidic-(HhH)_2_ with spermine caused an increase in the ^1^H chemical shift dispersion (**Figure 4**). The observation of new ^1^H signals upon the addition of spermine at >8.7 ppm, < 0.6 ppm, and between 7.3-7.6 ppm (**Figure 4**, peaks highlighted in grey), is in agreement with folding upon spermine binding. The appearance of methyl resonances at chemical shifts outside the “random coil” range (**Figure 4B**), and the increased dispersion of amide proton resonances in particular (**Figure 4A**), indicate a change in the chemical environment of these nuclei, most likely due to changes in structure. Chemical shifts of backbone protons tend to be shifted upfield for α-helices, by an average of −0.35 ppm relative to random coil values, versus downfield for β-sheets, by an average of +0.40 (38, 39).

Thus, the lack of new signals upon spermine addition in the 4.85-5.90 ppm region is consistent with the absence of β-sheet structure, while the observation of ^1^H signals at 3.8-4.4 ppm supports the presence of a α-helical structure (**Figure S2**). In the so-called “fingerprint region” of the 2D TOCSY spectrum (marked box, **Figure S2**) one would expect to observe one signal per residue for all non-proline residues (excluding the N-terminal residue). However, the presence of 21 glutamic acid residues, a duplicated sequence, and some spectral broadening – perhaps due to transient associations between protein molecules at the high protein concentration used for this experiment and in the presence of spermine – led to high ^1^HN resonance overlap, thus precluding the assignment of the ^1^H NMR peaks with this unlabeled protein sample. Nevertheless, the NMR data qualitatively support the chemical chaperone effects of spermine, where, in agreement with the CD analysis above, the structure of Acidic-(HhH)_2_ becomes more ordered and adopts a predominantly α-helical conformation upon spermine addition.

### Other polyamines and dications mediate folding of Acidic-(HhH)_2_

To better understand the chemical chaperone properties of polyamines, we performed titrations with natural polyamines of different length, as well as a mono-amine (propylamine) (**Figure 5** and **Figure S3**). Propylamine and NaCl had similar effects on the folding of Acidic-(HhH)_2_, both exhibiting a folding midpoint concentration around at ~0.4 M and complete folding at ~1 M. At approximately molar concentrations of salt, both charge masking and excluded volume effects can be significant (40) (**Figure S4**). In contrast, the chemical chaperone effect of polyamines is achieved at significantly lower concentrations (**Figure 5**). The strength of the chemical chaperone effect scaled with the number of amines per molecule, with spermine (a tetra-amine) exhibiting the strongest chemical chaperone and putrescene (a di-amine) exhibiting the weakest effect. The transition between putrescine (di-amine) and spermidine (a tri-amine) resulted in the greatest change in the apparent folding midpoint with a ~30-fold increase in potency. Finally, although ~100-fold less potent than the most potent polyamine (spermine), the dications Mg^2+^ and Ca^2+^ were far more active than either NaCl or propylamine, and even slightly more active than putrescine.

## Discussion

Acidic-(HhH)_2_ is a model primordial protein with a highly acidic surface. The ability of polyamines to efficiently fold Acidic-(HhH)_2_ suggests a possible role for polyamines as a chemical chaperone early in protein evolution, before basic amino acids could be readily incorporated into proteins. We note that other solutions to the basic amino acid problem have been reported, including the use of ornithine as an alternative basic amino acid (26) or a reliance on high concentrations of salt to drive protein folding (41), an effect we observe here as well. Similarly, phosphate binding sites are generally enriched for basic amino acids in modern proteins; however in the most ancient protein lineages phosphate binding is mediated by backbone interactions at the N-termini of α-helices (42), a feature also seen in the (HhH)_2_ fold (43). Dications, and specifically Mg^2+^ and Ca^2+^, also comprise an alternative solution and, although they were far less potent than spermine, they induced folding of Acidic-(HhH)_2_ at approximately seawater concentrations (44). Indeed, the prebiotic soup, which may have some resemblance to modern seawater, may have contained multiple chemical species that support the folding of simple, hyper-acidic proteins and nucleic acids. Indeed, evolution in general, and protein evolution in particular, is renowned for leveraging multiple solutions to tackle the same challenge. One needs only consider the great wealth of unrelated proteases that adopt different folds and employ different nucleophiles (45). In our view, an abundance of accessible solutions, which are complementary rather than contradictory, may well be a pre-requisite for the emergence of life.

Although framed and tested from the perspective of an acidic, primordial protein, these results lead us to hypothesize that polyamine-induced folding could be retained in modern biology – especially since the chemical chaperone benefits are observed within a biologically-relevant concentration range, particularly for spermine (1-2 mM in mammalian cells (46)) and spermidine (6-7 mM in *E. coli* (8)), though it is unknown what fraction of molecules are bound. In mammalian cells, the concentrations of various polyamines oscillate in accordance with the circadian rhythm (7), suggesting a role for polyamines in signaling and regulation (9, 47). Polyamines have also been shown to regulate individual proteins, such as a viral serine kinase (48) and the circadian regulation protein PER2 (7). However, the mechanistic and structural basis of these dependencies is unknown. Finally, we note that the high concentration of polyamines in bacteria is compatible with, and potentially related to, their generally acidic proteomes (49) – and hence, polyamines may have proteome-wide effects. We demonstrate that, mechanistically, polyamines can operate at concentrations with negligible ionic strength or excluded volume effects. Hence, specific binding interactions between structurally adjacent acidic residues are likely to be responsible for the chemical chaperone effects. Protein regions with high α-helicity that are enriched for acidic amino acids may therefore be a hallmark of a polyamine-responsive protein element.

Further studies are needed to support the above hypotheses, and also to examine whether hyper-acidic proteins could not only fold in the presence of polyamines, as demonstrated here, but also exert function. We could not observe dsDNA binding by Acidic-(HhH)_2_ in the presence of polyamines (data not shown), but we note that such measurements are technically challenging because polyamines bind nucleic acids on their own and the functional output of highly simplified proteins is often weak. Nonetheless, other model proteins may provide indications for function, thus lending further support for a role of polyamines at the earliest stages of protein evolution.

## Acknowledgements

This work was funded by the Israel Science Foundation grant 980/14 awarded to D.S.T. D.S.T. is the Nella and Leon Benoziyo Professor of Biochemistry. I.G.S acknowledges support by the Croatian Science Foundation (grant IP-2016-06-6272). T.S. is the incumbent of the Monroy-Marks Research Fellow Chair.

## Author contributions

I.G.S. proposed the idea of polyamines as substitutes to basic amino acids in the early proteins; D.D., L.M.L. and D.S.T. designed the experiments; D.D., L.M.L., E.A., A.K. performed protein production, CD and other biophysical experiments, and T.S. performed NMR measurements; D.D., L.M.L. and D.S.T. analyzed data; D.D., L.M.L. and D.S.T. wrote the paper.

