## Supplementary Information for "Polyamines Mediate Folding of Primordial Hyper-Acidic Helical Proteins"

### Supplementary Figures

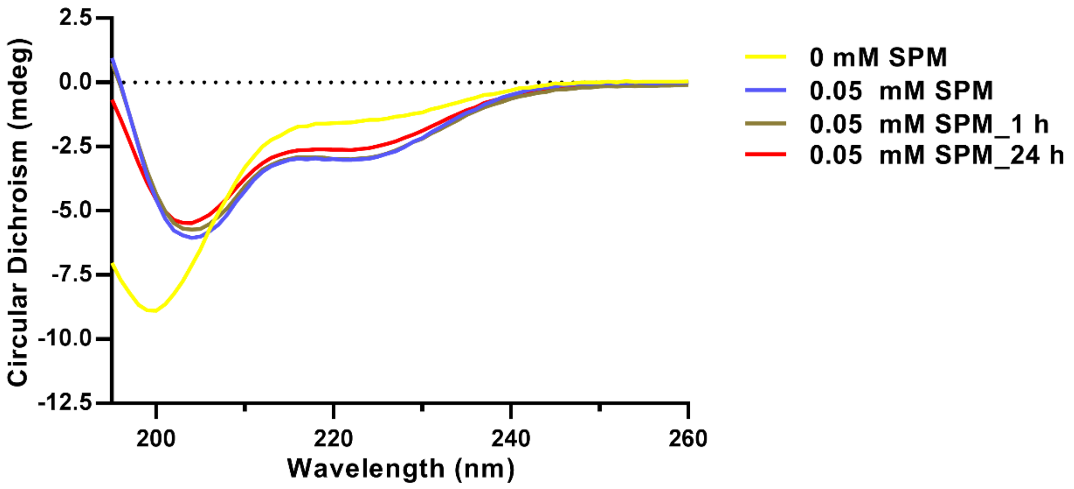


**Figure S1. Circular dichroism spectra of Acidic-(HhH)$_2$ upon incubation with spermine (SPM).** Spectra were taken immediately after addition of 0.05 mM spermine and again after 1 h and 24 h of incubation. The resulting spctrea are very similar, demonstrating that for Acidic-(HhH)$_2$ polyamine-induced structure formation is very fast.

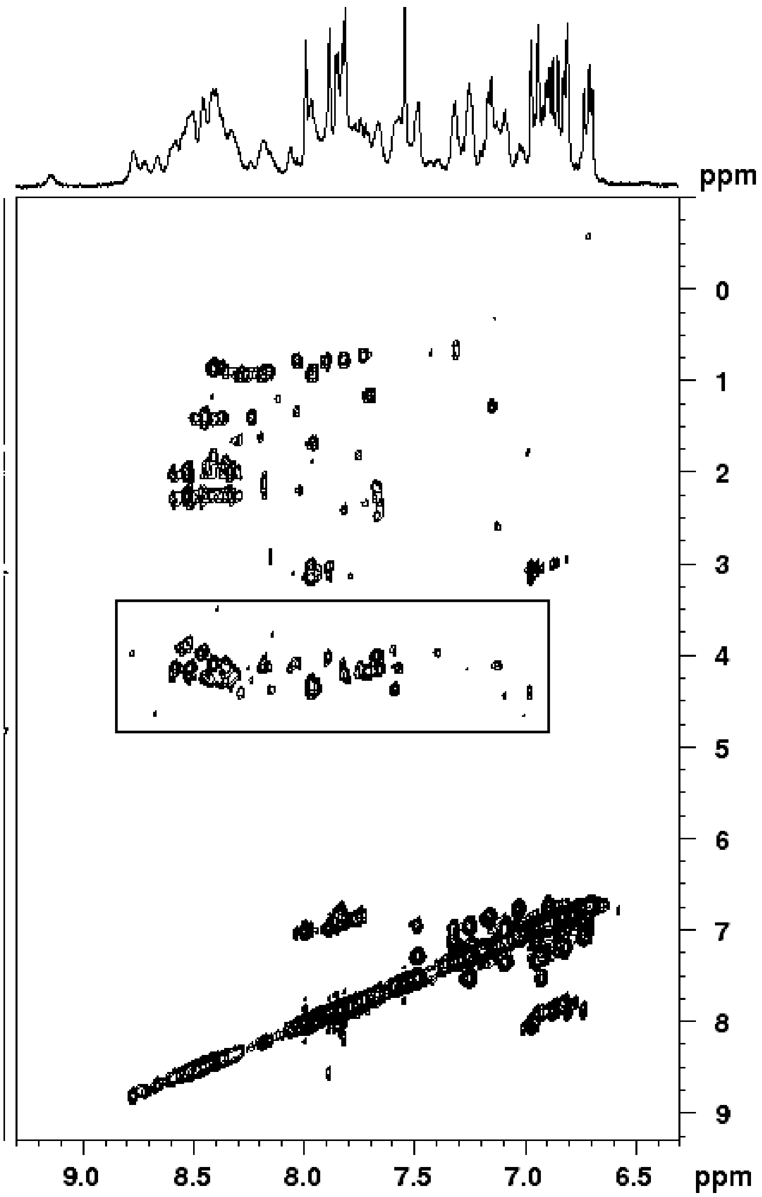


**Figure S2. A 2D $^{1}$H NMR TOCSY spectrum of Acidic-(HhH)$_2$ in the presence of 250 fold excess spermine at 293 K**. The spectrum was acquired on a 600 MHz NMR spectrometer using 120-ms mixing time and reports on intra-residue interactions of amide groups with aliphatic groups as well as interactions within aromatic sidechains. The fingerprints region of the spectrum, revealing intra-residue $^{1}$HN-$^{1}$H$^{\alpha}$ correlations, is marked with a box (see the main text).

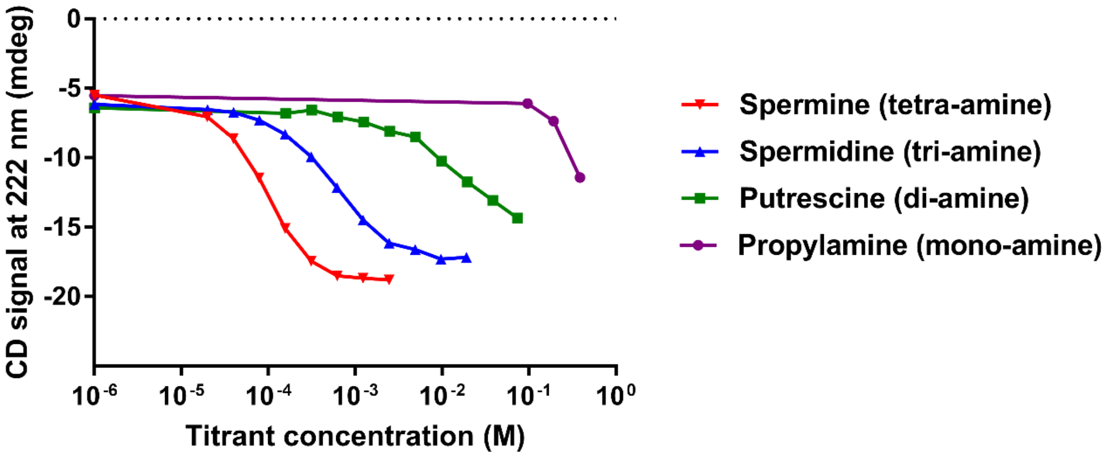


**Figure S3: Titration of Acidic-(HhH)$_2$ with various polyamines.** Circular dichroism spectra of 10 µM Ni-NTA-purified Acidic-(HhH)$_2$ were collected upon addition of various concentrations of polyamines. Plotted here is the CD signal at 222 nm, a reporter of α-helical structure, after buffer subtraction and dilution correction. Estimated midpoint concentrations are: spermine = 0.09 mM, spermidine = 0.6 mM, putrescine = 23 mM; these values and the midpoints concentrations from the independent titration in **Figure 5** are within ±20% (the midpoint for propylamine could not be reliably estimated from this dataset, but the plot is qualitatively very similar to the titration in **Figure 5**). Midpoint values were estimated from a linear interpolation between points and assuming a saturated folded signal of -19 mdeg at 222 nm.

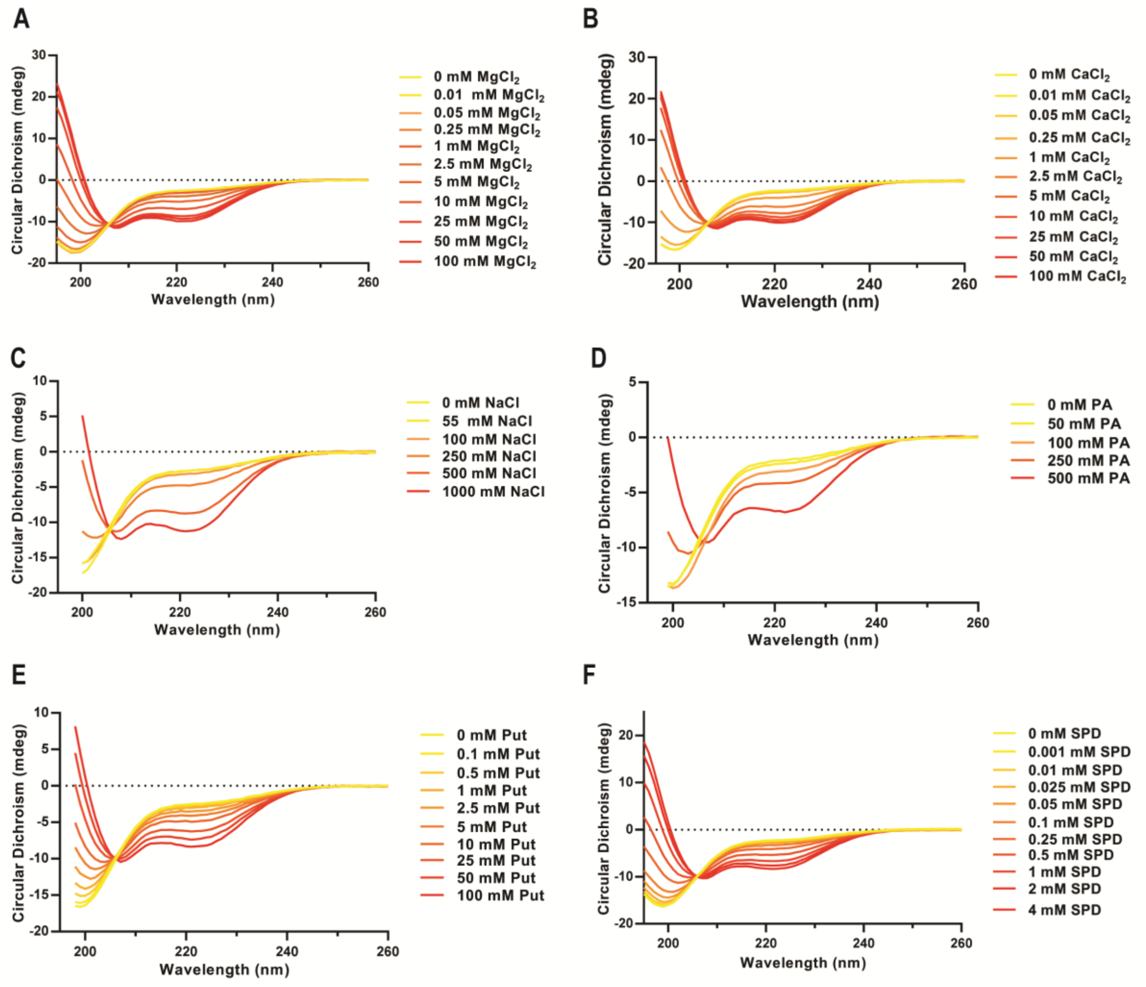


**Figure S4. Circular dichroism spectra of Acidic-(HhH)$_2$ titrated with salts and polyamines.** Circular dichroism spectra of 5 µM Acidic-(HhH)$_2$ were monitored upon addition of: **A**, $MgCl_2$; **B**, $CaCl_2$; **C**, NaCl; **D**, propylamine (PA); **E**, putrescine (Put); **F**, spermidine (SPD). Spectra from the titration with spermine are presented in **Figure 3**. Each curve represents the average of two spectra after buffer subtraction and correction for dilution due to titration. Data points exceeding 700 V of applied voltage to the photomultiplier tube (PMT) were discarded. These data were used to generate the titration curves presented in **Figure 5**.
